# Sex-dependent parameters of social behavior show marked variations between distinct laboratory mouse strains and their mixed offspring

**DOI:** 10.1101/2021.09.12.459932

**Authors:** Natalia Kopachev, Shai Netser, Shlomo Wagner

## Abstract

**Background:** The survival of individuals of gregarious species depends on their ability to properly form social interactions. In humans, atypical social behavior is a hallmark of several psychopathological conditions, such as depression and autism spectrum disorder, many of which have sex-specific manifestations. Various strains of laboratory mice are used to reveal the mechanisms mediating typical and atypical social behavior in mammals.

**Methods:** Here we used three social discrimination tests (social preference, social novelty preference, and sex preference) to characterize social behavior in males and females of three widely used laboratory mouse strains (C57BL/6J, BALB/c, and ICR).

**Results:** We found marked sex- and strain-specific differences in the preference exhibited by subjects in a test-dependent manner. Interestingly, we found some characteristics that were strain-dependent, while others were sex-dependent. Moreover, even in the social preference test, where both sexes of all strains prefer social over object stimuli, we revealed sex- and strain-specific differences in the behavioral dynamics. We then cross-bred C57BL/6J and BALB/c mice and demonstrated that the offspring of such cross-breeding exhibit a profile of social behavior which is different from both parental strains and depends on the specific combination of parental strains.

**Conclusions:** We conclude that social behavior of laboratory mice is highly sex- and strain-specific and strongly depends on genetic factors.

## 1. Introduction

The survival and success of individuals of gregarious mammalian species depend on their ability to form social interactions properly (1, 2). In humans, atypical social behavior is a hallmark of several psychopathological conditions and neurodevelopmental diseases (NDDs) (3), such as social anxiety disorder (4), autism spectrum disorder (5), and schizophrenia (6). Notably, many of these conditions have gender-specific manifestations and exhibit a robust diagnostic gender-bias (7, 8). However, unraveling the biological basis of gender-specific manifestations in Human NDDs is extremely difficult due to the strong influence of culture, education, and living style, all of which are heavily gender-biased (9). One way to overcome this difficulty is by using animal models, which are not affected by cultural factors (8, 10). Animal models are an important tool for exploring the biological basis of social behavior in general and particularly are used to unravel impaired mechanisms which underlie atypical social behavior in NDDs (11-14). Specifically, genetically modified mouse models carrying mutations in NDD-associated genes are widely used for such research (15). Notably, different laboratories use distinct laboratory mouse strains, some of which are inbred strains (16), for their research, and these strains also serve as a genetic background for the various mouse models of NDDs (17, 18). While multiple previous studies have explored inter-strain differences in murine social behavior (19-21), the effect of sex on aspects of social behaviors which are not associated with aggression, parenting, or sexual behavior has been poorly studied. Moreover, nothing is known about the consequences of mixing distinct strains by cross-breeding, regarding the social behavior of the offspring.

Several behavioral paradigms have been developed for assessing social behavior in mouse models. Of these, the most famous is the three-chamber test (22), which employs two types of social discrimination tasks: social preference (SP) and social novelty preference (SNP). We have previously presented a novel experimental system for automatic assessment of murine social discrimination behavior (23) and used it to analyze the behavior of C57BL/6J mice in the SP and SNP paradigms (21, 24). Here we added a third paradigm of social discrimination, the sex-preference (SxP) paradigm (25), in order to systematically examine sex- and strain-dependent behavioral parameters using three distinct strains of laboratory mouse strains (C57BL/6J, BALB/c, and ICR). Moreover, we characterized the behavior of offspring generated by cross-breeding of C57BL/6J and BALB/c mice.

## 2. Results

### 2.1 Sex- and strain-dependent differences in social discrimination behavior

To assess sex- and strain-specific differences in social behavior, we employed our computerized behavioral system (23, 24) to analyze the level and dynamics of investigation behavior exhibited by mice in three distinct social discrimination tests: SP, SNP, and SxP. We used all three tests to systematically characterize the behavior of male and female subjects of three distinct laboratory strains: C57BL/6J, BALB/c, and ICR (CD-1). Of these strains, the former two are inbred while the latter is an outbred strain. In all tests, we automatically measured the time dedicated by the subject to investigating each of two stimuli, simultaneously presented in distinct chambers, which are located at the opposite corners of the experimental arena. A statistically significant difference in investigation time between the two stimuli was considered to reflect a preference for the stimulus investigated by the subject for a longer duration. Notably, all experiments were repeated at least twice using distinct cohorts of animals, and the minimum number of mice subjected to each test was nineteen.

As apparent in Fig. 1, we found sex- and strain-specific differences in the performance of the subjects in a test-dependent manner. While a preference for the social stimulus compared to the object stimulus was observed for all strains and both sexes (Fig. 1a. *C57: Males: t*_*57*_*=7*.*754, p<0*.*0001; Females: t*_*26*_*=4*.*044, p<0*.*0001. BALB/c: Males: t*_*20*_*=10*.*210, p<0*.*0001; Females: t*_*18*_*=3*.*080, p=0*.*006. ICR: Males: t*_*22*_*=9*.*496, p<0*.*0001; Females: t*_*34*_*=6*.*110, p<0*.*0001, paired t-test*), the results of the other two tests were different. A preference for the novel social stimulus over the familiar one in the SNP test was observed for females of all strains. However, when examining males, only C57BL/6J mice showed such a preference (Fig. 1b. *C57: Males: t*_*57*_*=5*.*780, p<0*.*0001; Females: t*_*24*_*=3*.*088, p<0*.*005. BALB/c: Males: t*_*20*_*=1*.*423, n*.*s*.; *Females: t*_*19*_*=4*.*287, p<0*.*0001. ICR: Males: t*_*23*_*=1*.*199, n*.*s*.; *Females: t*_*34*_*=3*.*774, p<0*.*001, paired t-test*). The most variable pattern was found in the SxP test (Fig. 1c), where C57BL/6J and ICR males showed a clear preference for the opposite-sex stimulus, while BALB/c males did not show any preference between the stimuli (*C57: t*_*44*_*=4*.*281, p<0*.*0001; BALB/c: t*_*21*_*=1*.*812, n*.*s*.; *ICR: t*_*37*_*=9*.*508, p<0*.*0001, paired t-test*). In contrast, in none of the strains did females show a preference for the opposite-sex stimulus. However, ICR females showed a clear preference for the same-sex stimulus (*C57: t*_*49*_*=-0*.*982, n*.*s*.; *BALB/c: t*_*19*_*=0*.*739, n*.*s*.; *ICR: t*_*34*_*=5*.*833, p<0*.*0001, paired t-test*).

**Figure 1.**
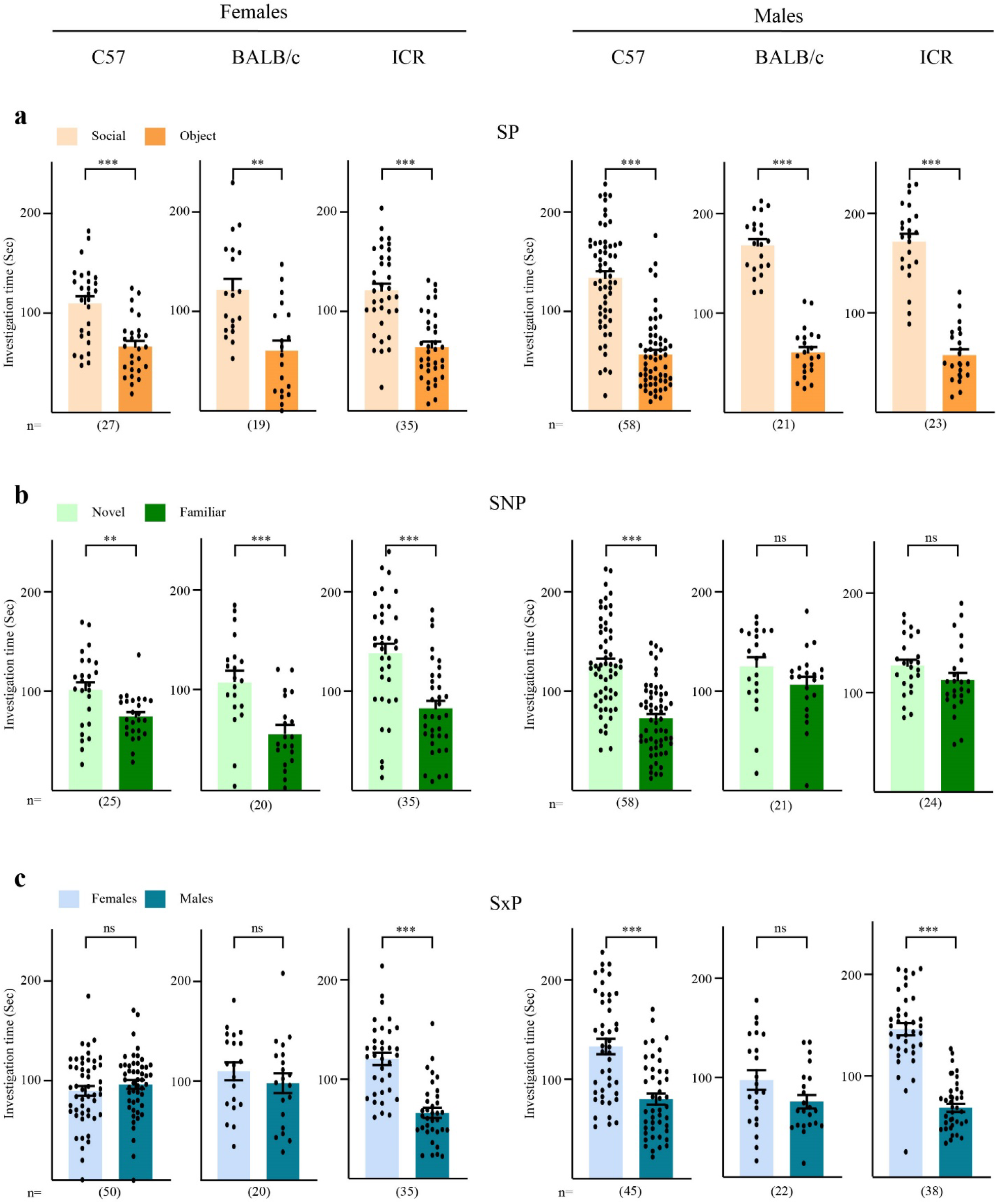
Sex- and strain-specific differences in social behavior during three distinct social discrimination tests: SP, SNP, and SxP. Mean investigation time (±SEM) measured separately for each stimulus across the SP (**a**), SNP (**b**), and SxP (**c**) tests, performed by females (left) and males (right) of all three mouse strains (denoted above). Sample size is denoted below the bars (\*\**p*<0.01, \*\*\**p*<0.001, ns – not significant, paired t-test).

For direct statistical comparison between animal groups, we calculated the difference in investigation time between the two stimuli (henceforth termed ΔIT) and compared it across sexes and strains separately for each test. As apparent in Fig. 2a, for the SP test, we found a significant main effect of the sex (*F*_*1,177*_*=19*.*241, p<0*.*0001, two-way ANOVA*). *Post hoc* analysis revealed a significantly higher ΔIT for males, as compared to females, for all three strains (*C57: t*_*83*_*=-2*.*085, p=0*.*040; BALB/c: t*_*38*_*=-2*.*113, p=0*.*041; ICR: t*_*56*_*=-3*.*787, p<0*.*0001, independent t-test*). In the SNP test (Fig. 2b), there was a significant interaction between sex and strain (*F*_*2,177*_*=4*.*864, p<0*.*009, two-way ANOVA*). *Post hoc* analysis revealed significantly higher ΔIT for female than male ICR mice (*t*_*57*_*=2*.*009, p<0*.*049*), while no significant difference was observed between male and female C57BL/6J and BALB/c mice (*C57: t*_*81*_*=-1*.*782, n*.*s*.; *BALB/c: t*_*39*_*=1*.*901, n*.*s*., *independent t-test*). Between strains, C57BL/6J male mice showed significantly higher ΔIT than ICR male mice (*p<0*.*040, Tukey’s HSD post hoc test*), while no significant difference was observed between females of the three strains. As for the SxP test (Fig. 2c), we observed again a significant interaction between sex and strain (*F*_*2,204*_*=12*.*705, p<0*.*0001, two-way ANOVA). Post hoc* analysis revealed significantly higher ΔIT for C57BL/6J and ICR males than their respective females (C57: *t*_*93*_*=-3*.*428, p<0*.*001; ICR: t*_*71*_*=-10*.*690, p<0*.*0001, independent t-test*). Across strains, significantly lower ΔIT values were found for ICR females compared to both C57BL/6J and BALB/c females (C57: p<0.001; BALB/c: *p<0*.*019, Tukey’s HSD post hoc test*), while ICR male mice showed significantly higher ΔIT than BALB/c male mice (*p<0*.*007, Tukey’s HSD post hoc test*). Overall, these data suggest sex- and strain-specific differences in the preference exhibited by subjects during the SP, SNP, and SxP tests, in a test-dependent manner.

**Figure 2.**
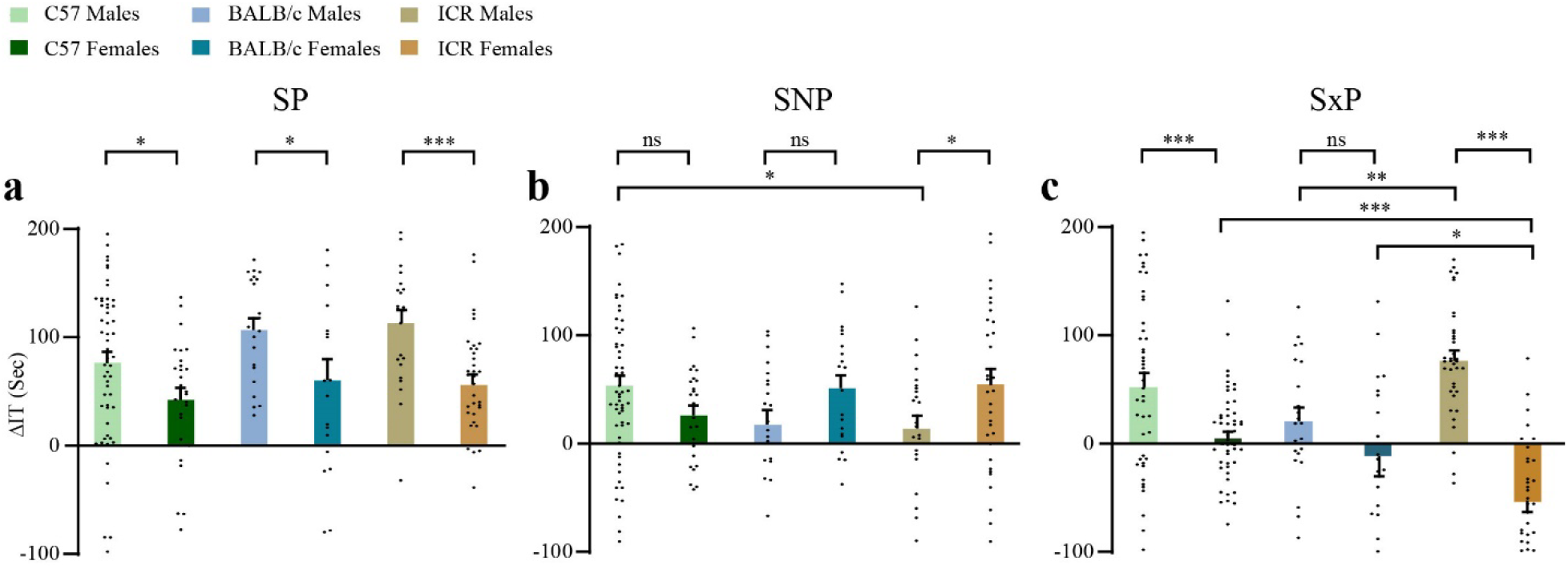
Sex- and strain-dependent difference in investigation time between the two stimuli in the SP, SNP, and SxP tests. Mean (±SEM) difference in investigation time (ΔIT) between the two stimuli, measured separately for each animal group during the SP (**a**), SNP (**b**), and SxP (**c**) tests. \**p*<0.05, \*\**p*<0.01, \*\*\**p*<0.001, ns – not significant, *post hoc* unpaired t-test or Tukey’s HSD test, following main effect in two-way ANOVA test.

### 2.2 Strain-, but not sex-dependent differences in the dynamics of transitions between stimuli

In a previous study, we found that C57BL/6J mice and Sprague-Dawley (SD) rats exhibit distinct dynamics of transitions between the two stimuli during the SP test and linked these differences to their distinct dynamics of social motivation (21). Therefore, in the current study we investigated the dynamics of transitions during all three tests for all animal groups. To avoid bias that may affect the behavioral dynamics towards the end of the experiment, we calculated the transitions that took place during the first four minutes of each trial. Interestingly, unlike the differences in investigation time (Fig. 1), we found no difference between males and females of any of the strains in the dynamics of transitions between stimuli (Fig. 3). We observed, however, marked differences among the various strains in a test-dependent manner. This difference was most notable between C57BL/6J mice and BALB/c mice. In accordance with our previous report, both male and female C57BL/6J mice exhibited high levels of transitions at the beginning of the test, which gradually declined during later stages. In contrast, BALB/c mice exhibited a rather constant pattern of transitions, except for the SxP test, were a lower level at the beginning of the test was observed among females. For ICR mice, the pattern of transitions was rather constant across the various tests with one exception: in the SxP test both male and female ICR mice exhibited a pattern of transitions which was similar to the one displayed by C57BL/6J mice.

**Figure 3.**
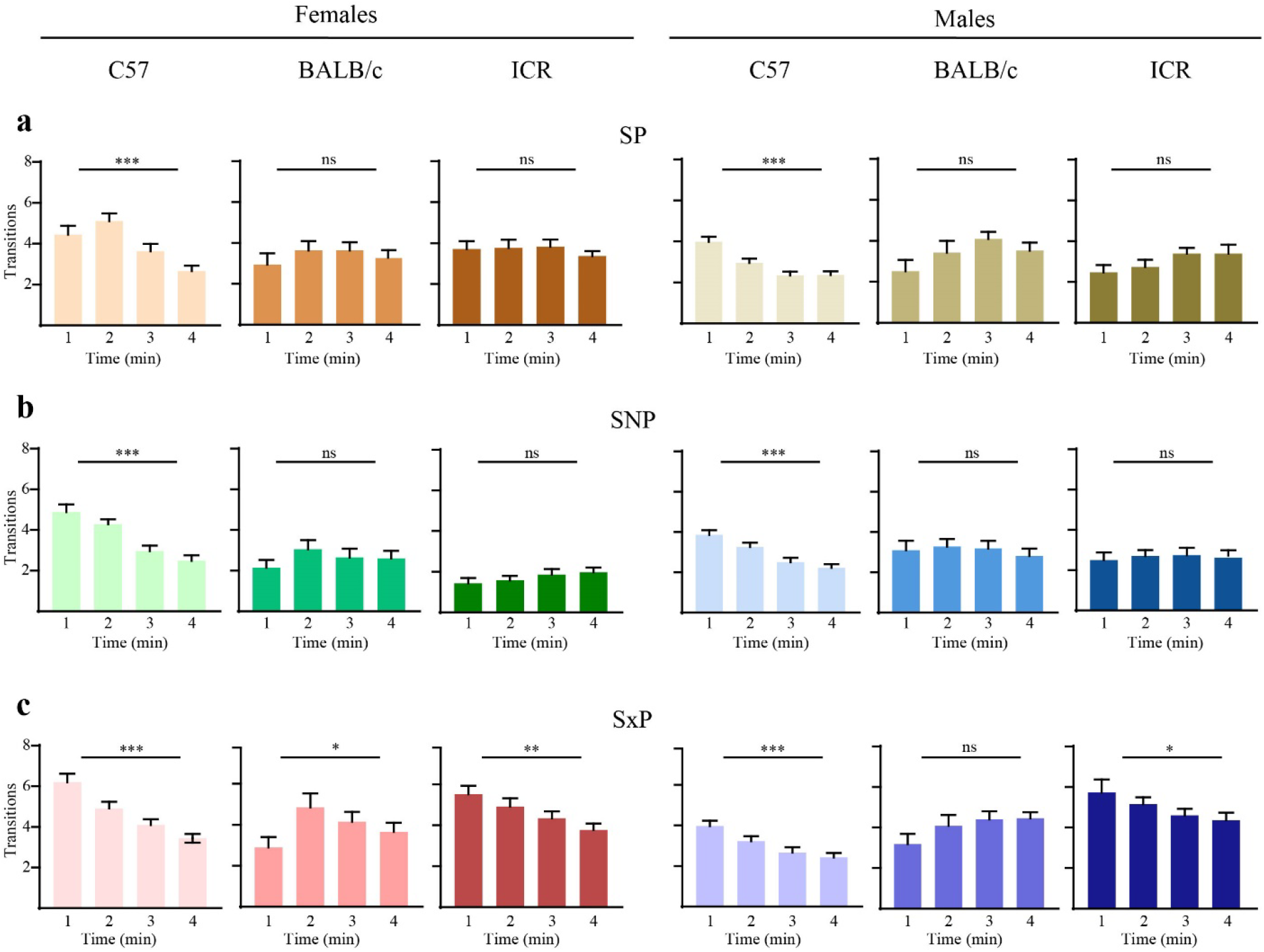
The dynamics of transitions between stimuli across the three behavioral tests. **a**. Mean number of transitions made by male (left) and female (right) subjects of the distinct mouse strains (denoted above) during the SP test (1-min bins). **b**. As in **a**, for the SNP test. **c**. As in **a**, for the SxP test. \**p*<0.05,\*\**p*<0.1, \*\*\**p*<0.01, one-way repeated ANOVA.

For statistical comparison between the groups, we calculated the difference in transition number between the first and fourth minute of the test (henceforth termed ΔTransition), for each animal group. These time bins were selected since they displayed the greatest difference in transition number in most cases. For the SP test (Fig. 4a), we found a significant main effect of the strain but not sex (*Strain: F*_*2,177*_*=14*.*330, p<0*.*0001; Sex: F*_*1,177*_*=2*.*872, n*.*s*., *two-way ANOVA*). *Post hoc* analysis revealed a significant difference between C57BL/6J males and males of both other strains (*C57 vs. BALB/c: p<0*.*0001; C57 vs. ICR: p<0*.*0001, Tukey’s HSD post hoc test*), while C57BL/6J female mice showed a significant difference from BALB/c females only (*C57 vs. BALB/c: p<0*.*026, Tukey’s HSD post hoc test*). A slightly different pattern was found for the SNP test (Fig. 4b), with *post hoc* analysis following main effect of strain (*Strain: F*_*2,177*_*=19*.*962, p<0*.*0001, two-way ANOVA*) revealing significant differences between C57BL/6J female mice and females of both other strains (*C57 vs. BALB/c: p<0*.*0001; C57 vs. ICR: p<0*.*0001, Tukey’s HSD post hoc test*), while male C5BL/6J mice showed significant difference from ICR males only (*C57 vs. ICR: p<0*.*008, Tukey’s HSD post hoc test*). For the SxP test (Fig. 4c), *post hoc* analysis following main effect of strain (*Strain: F*_*2,204*_*=11*.*639, p<0*.*0001, two-way ANOVA*) revealed that C57BL/6J and ICR males differed from BALB/c males, while ICR females differed from BALB/c females (*Males: C57 vs. BALB/c: p<0*.*001; BALB/c vs. ICR: p<0*.*002; Females: BALB/c vs. ICR: p<0*.*011, Tukey’s HSD post hoc test*). Overall, these results suggest that the transition dynamics are a strain-, but not sex-specific feature, which is kept rather unchanged across the various tests, at least for the inbred C57BL/6j and BALB/c mice.

**Figure 4.**
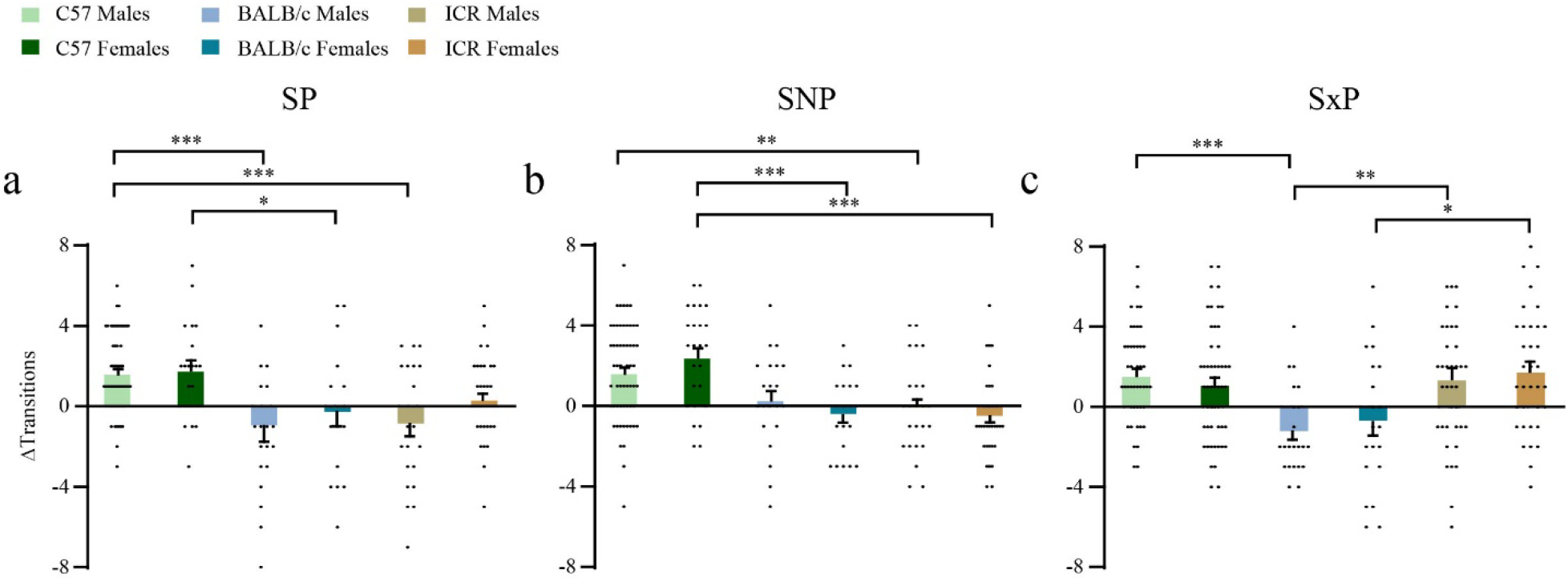
Sex-but not strain-dependent difference in the dynamics of transitions between the two stimuli. Mean (±SEM) difference in the number of transitions between the two stimuli between the first and fourth minute of each test (ΔTransition), measured separately for each animal group during the SP (**a**), SNP (**b**), and SxP (**c**) tests. \**p*<0.05, \*\**p*<0.01, \*\*\**p*<0.001, *post hoc* unpaired t-test or Tukey’s HSD test, following main effect in two-way ANOVA test.

### 2.3 Sex- and strain-dependent differences in the dynamics of social preference behavior

We next analyzed the dynamics of investigation behavior across the various strains and sexes. For this analysis, we focused on the SP test for two reasons. First, since all groups showed a significant preference of the social stimulus over the object, we reasoned that differences in behavioral dynamics during this test could not be attributed to preference variations. Second, as this test was always the first to be performed, it could not be affected by other tests in a strain- or sex-dependent manner. We observed an apparent difference between males and females in the behavioral dynamics of the SP test (Fig. 5a). While males of all strains showed social preference that was very strong at the beginning of the test and declined over time, females showed a relatively stable preference throughout the test. To statistically analyze these apparent differences, we compared ΔIT between the first and last two minutes of the test, across sex and strains (Fig. 5b). We found an interaction between time and sex (*F*_*1,177*_*=16*.*468, p=0*.*0001, mixed-model ANOVA*). *Post hoc* analysis revealed a significant difference between the first and last two minutes only for males, but not for females of all strains (*C57: Males: t*_*57*_*=7*.*017, p=0*.*0001; Females: t*_*26*_*=1*.*182, n*.*s*., *BALB/c: Males: t*_*20*_*=2*.*242, p=0*.*036; Females: t*_*18*_*=-1*.*170, n*.*s*., *ICR: Males: t*_*22*_*=3*.*079, p<0*.*005; Females: t*_*34*_*=-0*.*022, n*.*s*., *paired t-test*). Thus, the behavioral dynamics during the SP test were sex-dependent.

**Figure 5.**
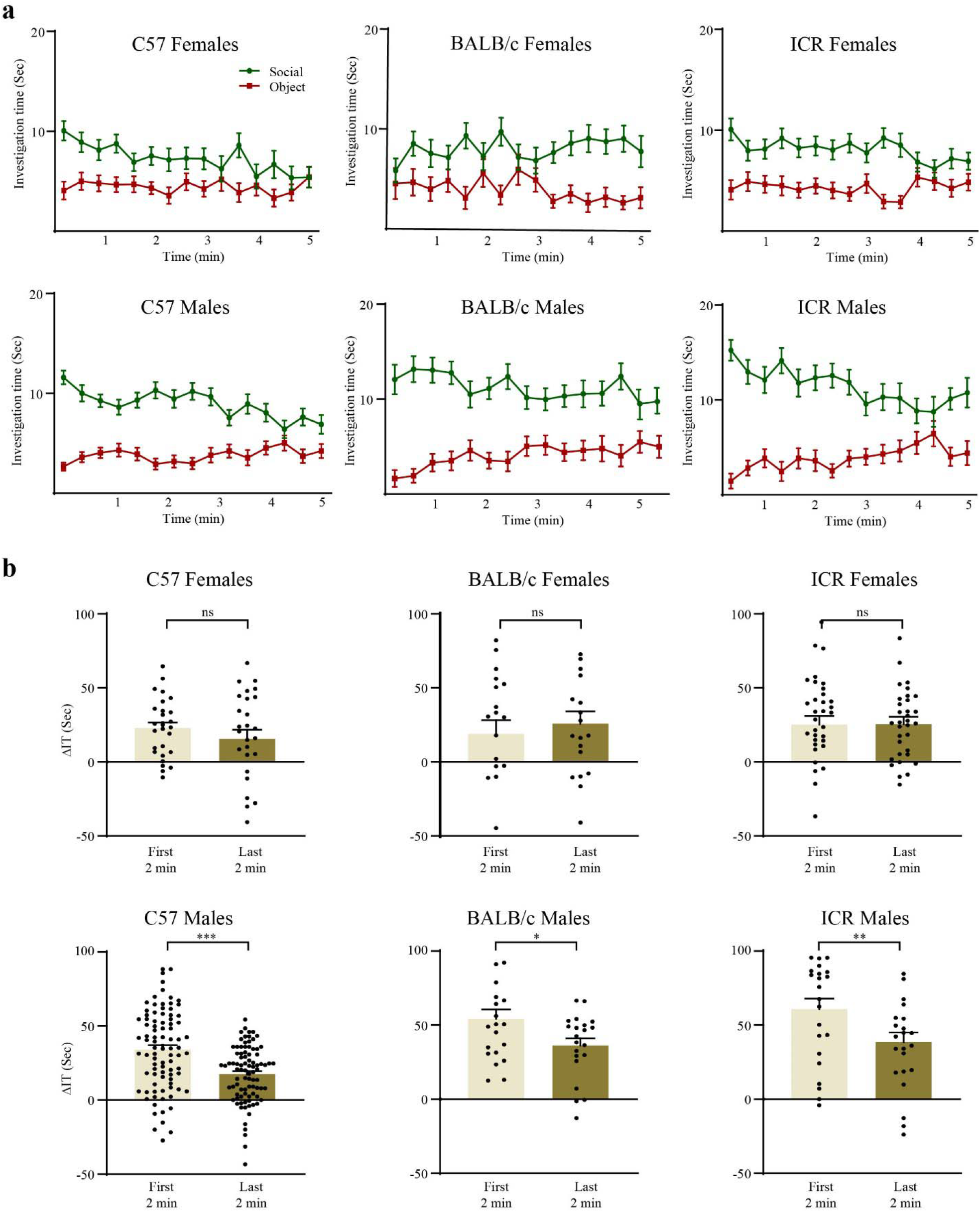
Sex-dependent dynamics of investigation behavior during the SP test. **(a)** Mean investigation time (±SEM), measured separately for each stimulus (20-s bins) along the time course of the SP test session (20-s bins) for females (upper panels) and males (lower panels) of the three strains. Note the transiently strong preference of males as compared to the stable but weaker preference of females. **(b)** Mean ΔIT (±SEM) measured separately for the first and last two minutes of SP test. \**p*<0.05, \*\**p*<0.01, \*\*\**p*<0.001, ns – not significant, *post hoc* paired t-test following the main effect in mixed-model ANOVA test.

### 2.4. Sex-dependent differences in bout duration during the SP test

We have previously shown that in C57BL/6J mice the difference in investigation time between the social stimulus and the object during the SP test is reflected only in long (>6 s) investigation bouts, while shorter bouts (≤6 s) do not differ between the two stimuli (24). To examine this point across sexes and strains, we plotted the fraction of ΔIT as a function of investigation bout duration for all strains and compared it between males and females. As for the transitions, we calculated only the bouts that took place during the first four minutes of each trial, in order to avoid bias affecting the behavioral dynamics towards the end of the experiment. As apparent in Fig. 6a, in all strains there seems to be a clear difference between males and females. Males of all strains showed longer bouts than females. In addition, BALB/c and ICR females showed more short bouts than males. We then statistically compared ΔIT separately for short and long bouts across sexes and strains. For short bouts (Fig. 6b), we found a significant interaction between sex and strain (*F*_*2,177*_*=6*.*834, p<0*.*001, two-way ANOVA*), with a *post hoc* analysis showing a higher ΔIT for female BALB/c and ICR, but not C57BL/6J mice, as compared to males (*BALB/c: t*_*38*_*=4*.*286, p<0*.*0001; ICR: t*_*56*_*=2*.*211, p=0*.*039, independent t-test*). Also, BALB/c showed a significantly higher ΔIT as compared to C57BL/6J females, while no difference was found between the males (Females: *C57 vs. BALB/c: p<0*.*005; Tukey’s HSD post hoc test*). In contrast, long bouts (Fig. 6c) showed the main effect in ΔIT only for the sex (*F*_*1,177*_*=36*.*022, p<0*.*0001*), with a *post hoc* analysis revealing a significantly higher ΔIT for males as compared to females in all strains (*C57: t*_*83*_*=-2*.*130, p=0*.*036; BALB/c: t*_*38*_*=-3*.*916, p<0*.*0001; ICR: t*_*56*_*=-4*.*616, p<0*.*0001, independent t-test*). Thus, it seems as if social preference of males is generally expressed by longer bouts, while females express their social preference using shorter investigation bouts.

**Figure 6.**
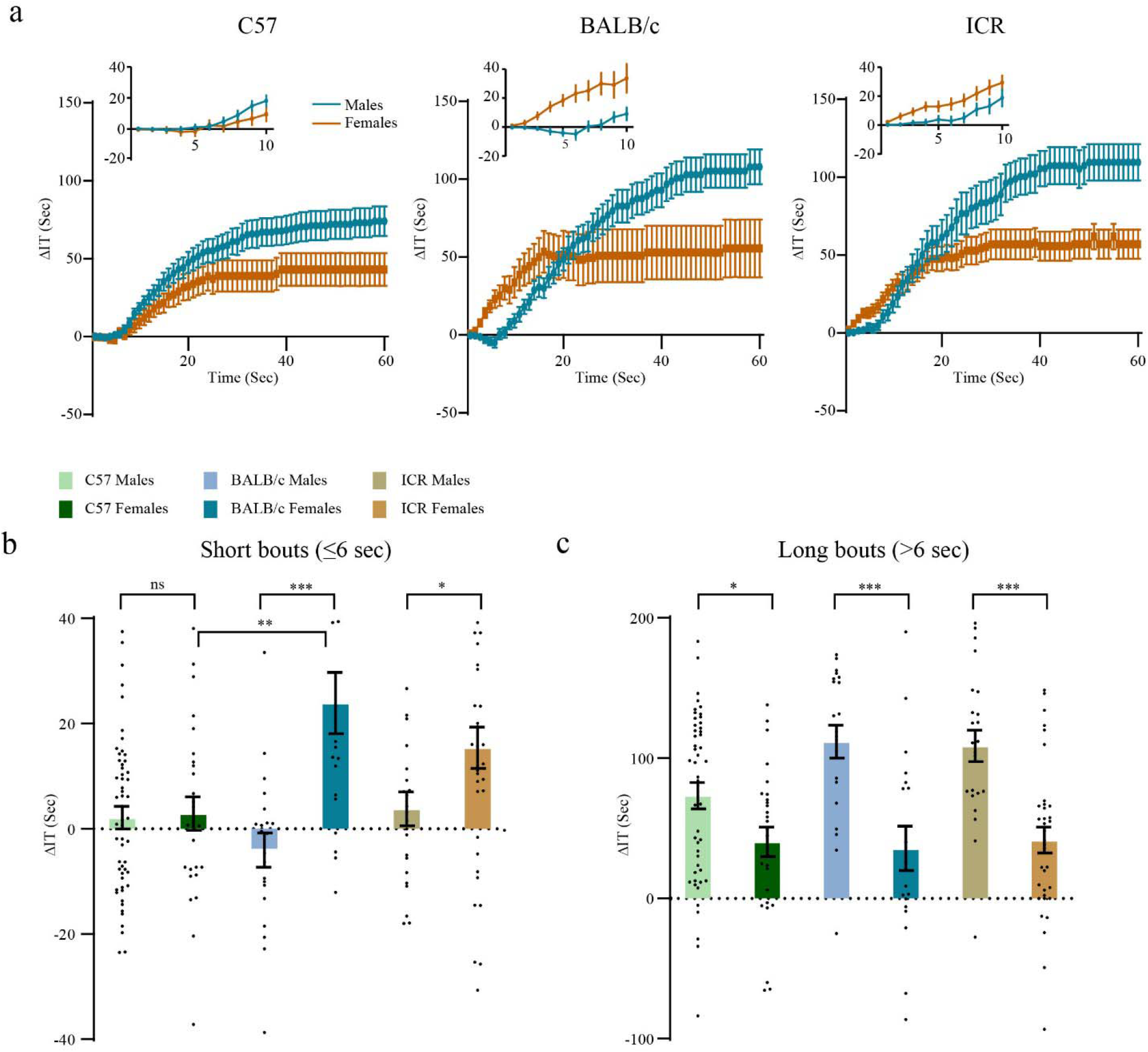
Sex-dependent differences in bout duration during the SP test. **(a)** Cumulative ΔIT plotted against bout duration during the SP test, separately for males (blue) and females (brown) of the three strains. Inset – same data on a larger scale for duration<10 s. **(b)** Mean ΔIT (±SEM), calculated separately for short (≤6 s) investigation bouts generated during the SP test by each animal group. **(c)** As in B, for long (>6 s) bouts. \**p*<0.05, \*\**p*<0.01, \*\*\**p*<0.001, ns – not significant, *post hoc* unpaired t-test or Tukey’s HSD test, following main effect in two-way ANOVA test.

### 2.5. Offspring of a cross-breeding between C57BL/6J and BALB/c mice exhibit distinct social behavioral characteristics which depend on their parental combination

Finally, as we found significant differences in social behavior between the distinct strains, we sought to explore the effect of cross-breeding two distinct strains. For this purpose, we focused on male offspring generated by crossing C57BL/6J and BALB/c mice, both of which are inbred strains of a similar size. We separately analyzed litters of C57BL/6J mothers and BALB/c fathers, termed by us CB mice, and litters of BALB/c mothers and C57BL/6J fathers, termed BC mice. Interestingly, we found that the BC and CB offspring showed different behavioral profiles, neither of which recapitulated the profiles of any of their parental strains. In the SP test (Fig. 7a), we found that only CB mice showed a significant social preference (*BC: t*_*35*_*=-0*.*298, n*.*s*.; *CB: t*_*46*_*=2*.*903, p=0*.*006, paired t-test*). Nonetheless, the two cross-bred strains were similar to each other in the dynamics of their behavior, with an initial preference at the beginning of the test and a loss of preference at a later stage (Fig. 7b), a pattern not observed in any of the parental strains (Fig. 5a). As for the transitions, both groups showed a pattern resembling their mothers, with BC mice showing a pattern similar to BALB/c mice and CB mice to C57BL/6J mice (Fig. 7c). In the SNP test (Fig. 7d), both BC and CB mice did not show any preference, similar to BALB/c males but in contrast to C57BL/6J mice (*BC: t*_*18*_*=0*.*531, n*.*s*.; *CB: t*_*23*_*=1*.*137, n*.*s*., *paired t-test*). The behavioral similarity of both groups to BALB/c mice was also apparent in the dynamics of the investigation behavior (Fig. 7e) and transitions (Fig. 7f). In the case of the SxP test, while both CB and BC mice exhibited a significant preference for the female over the male stimulus (Fig. 7g), this preference looked stronger in CB mice, similar to C57BL/6J males, and much weaker in BC mice, similar to BALB/c males, which did not show any sex preference (*BC: t*_*32*_*=3*.*430, p<0*.*002; CB: t*_*44*_*=12*.*926, p<0*.*0001, paired t-test*). These differences were also apparent from the dynamics of investigation behavior (Fig. 7h) and transitions (Fig. 7i). Thus, in the SxP test, both types of cross-bred offspring showed a behavioral pattern which resembles males of their maternal strain.

**Figure 7.**
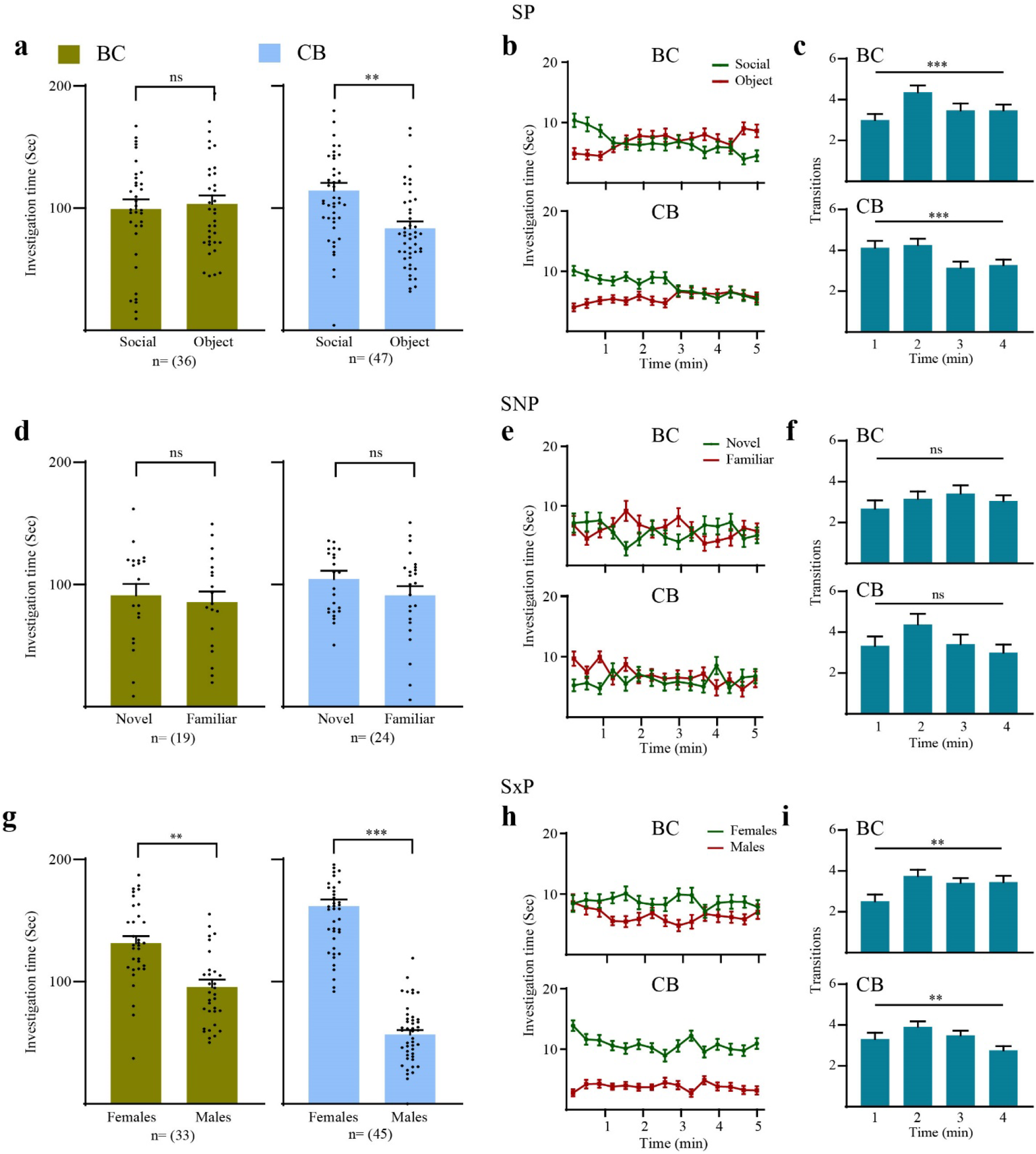
Offspring of a cross-breeding between C57BL/6J and BALB/c mice exhibit distinct social behavioral characteristic. **(a)** Mean investigation time for each stimulus during the SP test for BC (green) and CB (blue) mice. **p<0.01, paired t-test. **(b)** Mean investigation time measured separately for each stimulus (20-s bins) along the time course of the SP test session (20-s bins) for BC (upper panel) and CB (lower panel) mice. **(c)** Mean number of transitions between the two stimuli made during the SP test for BC (upper panel) and CB (lower panel) subjects. \*\*\**p*<0.001, one-way repeated ANOVA. **(d-f)** As in **a-c**, for the SNP test. **(g)** As in **a**, for the SxP test. \*\**p*<0.01, \*\*\**p*<0.001, paired t-test. **(h)** As in **b**, for the SxP test. **(i)** As in **c**, for the SxP test. \*\**p*<0.01, one-way repeated ANOVA

For statistical comparison between the groups, we calculated the ΔIT (Fig. 8a) and ΔTransitions (Fig. 8b) and compared between BC and CB male mice separately for each test. For both parameters, we found that CB mice exhibited significantly higher values than BC mice in both the SP (ΔIT: *t*_*81*_*=-2*.*049, p<0*.*044;* ΔTransitions: *t*_*81*_*=-2*.*672, p<0*.*009, independent t-test*) and SxP (ΔIT: *t*_*76*_*=-5*.*308, p<0*.*0001;* ΔTransitions: *t*_*76*_*=-2*.*944, p<0*.*004, independent t-test*) tests, while no differences were found in the SNP test (ΔIT: *t*_*41*_*=-0*.*466, n*.*s*.; ΔTransitions: *t*_*41*_*=-1*.*041, n*.*s*., *independent t-test*). Notably, although the difference in ΔTransitions was not significant in the SNP test, the trend was similar to the other two tests. Thus, regarding the dynamics of transitions, it seems as if in all cases the offspring behavior resembled their mothers’ behaviors. We conclude that offspring of BALB/c and C57BL/6J mice exhibit a profile of social behavior which depends on the parental combination and is different from both parental strains, with a tendency to mimic the behavior of the maternal strain.

**Figure 8.**
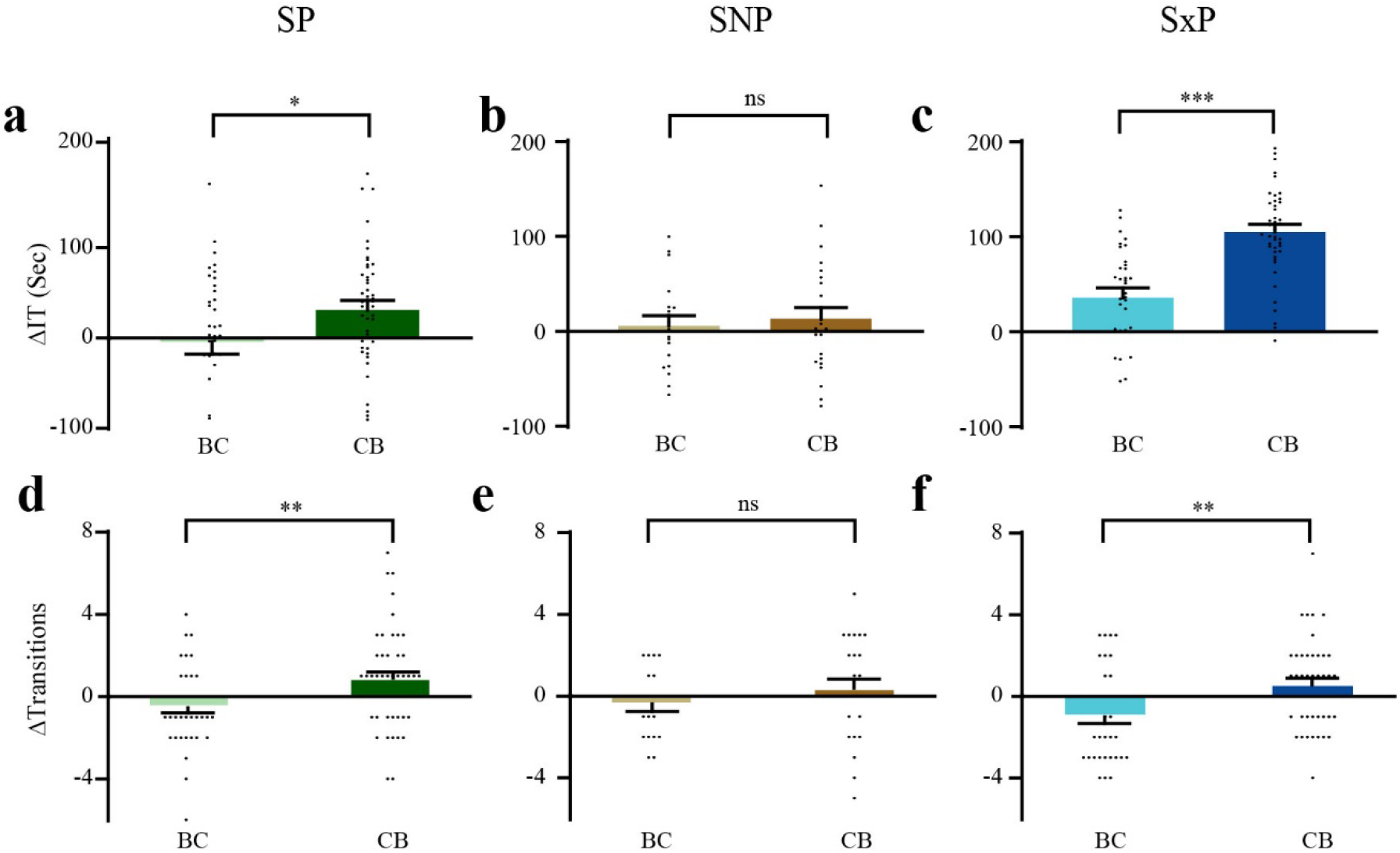
Comparing offspring of C57BL/6J mothers and BALB/c fathers with offspring of BALB/c mothers and C57BL/6J fathers. **(a-c)** Mean ΔIT, for the SP (**a**), SNP (**b**), and SxP (**c**) results of BC and CB mice, shown in Fig. 7**a, d, g**, respectively. (**d-f**) Mean ΔTransitions, for the SP (**d**), SNP (**e**), and SxP (**f**) results of BC and CB mice, shown in Fig. 7**c, f, i**, respectively. \**p*<0.01, \*\**p*<0.01, \*\*\**p*<0.001, ns – not significant, unpaired t-test.

## 3. Discussion

Sex-dependent differences in murine social behavior have been comprehensively explored in previous studies, but this was done mainly regarding hormonal-driven aspects of social behavior (26, 27), such as sexual (28), anxiety/aggressive (10), and parental behaviors (29). In contrast, evidence for differences in other aspects, such as social preference and social novelty preference have been reported only anecdotally, and in some cases with contradicting results (see for example 22, 30). Here, we systematically explored such differences across three types of social discrimination tasks and three distinct laboratory mouse strains. We have used a relatively large number of animals and at least two independent cohorts for each animal group, to verify replicability. We found a marked difference in social investigation behavior between males and female for all three strains, especially in the SNP and SxP tests. While the SxP test may be related to sexual behavior (31), the SNP test is not, and is well known to reflect social novelty seeking. Thus, our results demonstrate that even social behaviors which are not directly related to the sexual, aggressive, or parental aspects may be sex-dependent.

Our computerized experimental system enables analyzing the dynamics of the social discrimination behavior as well as categorizing each investigation bout according to its duration (23, 24). Using these features, we found that even in the SP test, where all strains and sexes exhibit a significant preference for a social stimulus over an object, the dynamics of the behavior may markedly differ between strains and sexes. This was most profoundly demonstrated by the difference between ICR and BALB/c males, which exhibited a significant change over time in their social preference behavior, and all other groups. We have previously shown that this reduction over time in social preference reflects dynamic changes in the motivation for social interactions. Thus, ICR and BALB/c males seem to exhibit a very strong drive for social interactions at the beginning of the encounter, which gradually decreases over time, while the other groups show a lower level of social motivation, which is kept rather constant over time.

Another dynamic aspect of social preference found to be sex-specific is the duration of investigation bouts. We have previously shown for male C57BL/6J mice that there is no difference in investigation using short bouts between the stimuli. In contrast, the preference for the social stimulus is specifically expressed by more investigation using long bouts. We thus suggested that long bouts reflect the interaction of the subject with the social stimulus while short bouts reflect curiosity per se. Here, we show for the first time that in females, specifically in ICR and BALB/c females, the picture is quite different. In contrast to males, females express their social preference by both long and short bouts, with no apparent difference between them. Moreover, in all strains, males exhibited longer investigation bouts towards the social stimulus than females. This sex-specific difference in the duration of investigation bouts may reflect a weaker motivation for interaction with a novel same-sex social stimulus exhibited by females, compared to males.

Another aspect of the behavioral dynamics we explored is the transitions between the two stimuli. Interestingly, while most differences in investigation behavior were between males and females, we found no sex-dependent differences in the pattern of transitions. Instead, this characteristic seems highly strain-dependent. For example, the C57BL/6J and BALB/c strains exhibited a rather uniform and specific pattern across both sexes and all tests, while ICR mice showed a pattern similar to BALB/c mice in the SP and SNP tests and a pattern similar to C57BL/6J mice in the SxP test. In a previous study (21) we suggested that similarly different patterns of transitions between SD rats and C57BL/6J mice reflect their distinct dynamics of social motivation. Accordingly, a recent study found a specifically high level of activity in dopaminergic neurons of the ventral tegmental area (VTA) during transition from object to social stimuli (30). Thus, the strain-dependent distinct patterns of transitions found by us may reflect distinct dynamics of social motivation between the strains.

The genetic basis of social behavior is well established throughout the animal kingdom. Significant differences between various mouse strains have been previously found using the three-chamber test (19, 20, 22). Yet, to our knowledge, our study is the first to examine the consequences of cross-breeding between distinct mouse strains which exhibit different social behavior. Interestingly, the behavior of F1 male offspring of the cross-breeding scheme was different from both parental strains in a test-dependent manner. This was most strikingly demonstrated by the dynamics of the social preference behavior, where both BC and CB mice showed such preference only at the beginning of the test. Such a phenomenon, of F1 behavior which is weaker than both parental mouse strains, was previously shown for morphine analgesia (32). Even more surprising is the observation that F1 offspring that were born to male C57BL/6J mothers and BALB/c fathers, differ in their behavior compared to offspring of BALB/c mothers and C57BL/6J fathers. Interestingly, in several aspects of their behavior, the F1 offspring showed resemblance to the strain of their mothers. The question whether this difference between BC and CB mice is caused by genetic, epigenetic, or environmental (i.e., the strain of the female taking care of the newborn animals) factors should be addressed by future studies.

Overall, we conclude that social behavior of laboratory mice, even if not related to sexual, aggressive, or parental aspects, is highly sex-and strain dependent. Moreover, we show that the behavioral outcomes of a cross-breeding between mouse strains is unpredictable and may differ markedly between tests and breeding schemes. These conclusions should be taken into account in future studies exploring modified social behavior in genetic mouse models.

## 4. Materials and Methods

### Animals

The research subjects were naïve male and female adult (8-12 weeks old) mice of three distinct laboratory strains: C57BL/6J, BALB/c, and ICR (CD-1) and adult male mice generated by crossing C57BL/6J and BALB/c mice. The social stimuli were C57BL/6J, BALB/c, and ICR (CD-1) juvenile (21-30 day-old) naïve male and female mice (SP, SNP), and C57BL/6J, BALB/c, and ICR (CD-1) adult (8-12 weeks old) naïve male and female mice (SxP). Mice were commercially obtained (Envigo, Israel) and housed in Plexiglas cages in groups of 2-5 animals per cage. They were kept at 22±2°C under a 12-h light/12-h dark cycle, with lights being turned on at 7 p.m. each night. All animals had *ad libitum* access to food (standard chow diet; Envigo RMS, Israel) and water. Behavioral experiments were performed during the dark phase, under dim red light. All experiments were performed according to the National Institutes of Health guide for the care and use of laboratory animals and approved by the Institutional Animal Care and Use Committee (IACUC) of the University of Haifa.

### Behavioral assays

All social discrimination tasks were conducted using our published automated experimental system (23). SP and SNP tests were conducted on the same day, as previously described (23). The SxP test was conducted as previously described (25). Briefly, it consisted of 15 min habituation to the arena with empty chambers, followed by exposing the subject for 5 min to both novel adult male and female social stimuli located in individual chambers at opposite corners of the arena. All stimuli used in all three tests (SP, SNP, and SxP) were C57BL/6J, BALB/c, and ICR (CD-1) mice.

### Data analysis

Video data analysis was conducted by our published custom-made TrackRodent software, as previously described in detail (23).

### Statistical analysis

All statistical tests were performed using SPSS 23 (IBM) statistics software. Shapiro-Wilk test was used for examining the normal distribution of the dependent variables. A 2-tailed paired t-test was used to compare between parameters within a group, and a 2-tailed independent t-test was used to compare a single parameter between distinct groups. For examining the influence of one categorical independent variable on one continuous dependent variable, a one-way ANOVA model was applied to the data. This model assesses the main effect of the independent variable on the dependent variable. For examining the influence of two different categorical independent variables on one continuous dependent variable, a two-way ANOVA model was applied to the data. This model assesses the main effect of each independent variable and the interaction between them. For comparison between multiple groups and parameters, a mixed-model analysis of variance (ANOVA) model was applied to the data. This model contains one random effect (ID), one within effect, one between effect, and the interaction between them. All ANOVA tests were followed, if the main effect or interaction were significant, by *post hoc* Student’s t-test with Bonferroni’s correction. Significance was set at p-value *<0.05.

## Supplementary Materials

None

## Author Contributions

Conceptualization, S.W. and S.N.; methodology, S.N.; software, S.N.; validation, S.W. and S.N.; formal analysis, S.N.; investigation, N.K.; resources, S.W.; data curation, S.N.; writing—original draft preparation, S.W; visualization, S.N and N.K..; supervision, S.W.; project administration, S. N.; funding acquisition, S.W. All authors have read and agreed to the published version of the manuscript.

## Institutional Review Board Statement

All experiments were performed according to the National Institutes of Health guide for the care and use of laboratory animals and approved by the Institutional Animal Care and Use Committee (IACUC) of the University of Haifa.

## Code availability

All codes and datasets used for the current study are available on request from the corresponding author within a reasonable time. The code used for the video analysis is publicly available at the following link: [https://github.com/shainetser/TrackRodent].

